# WaspAtlas: A *Nasonia vitripennis* gene database

**DOI:** 10.1101/020669

**Authors:** Nathaniel J. Davies, Eran Tauber

## Abstract

**Summary:** WaspAtlas is a new integrated gene database for the emerging model organism *Nasonia vitripennis*, which combines annotation data from all available annotation releases with original analyses to form the most comprehensive *N. vitripennis* resource to date. WaspAtlas allows users to browse and search for gene information in a clear and co-ordinated fashion providing detailed illustrations and easy to understand summaries. The database provides a platform for integrating gene expression and DNA methylation data. WaspAtlas also functions as an archive for empirical data relating to genes, allowing users to easily browse published data relating to their gene(s) of interest.

**Availability:** Freely available on the web at http://waspatlas.com. Website implemented in Catalyst, MySQL and Apache, with all major browsers supported.

## 1 INTRODUCTION

*Nasonia vitripennis* is a non-social parasitoid wasp which is becoming an important model organism in many areas of study. The emergence of *N. vitripennis* as a model organism is explained by the fact that it offers several important advantages over other insect models. Such advantages include but are not limited to a haplodiploid sex determination system (Werren and Loehlin 2009), simple rearing, a fully functional DNA methylation system (Beeler et al., 2014, Park et al., 2011, Wang et al., 2013), robust circadian (Bertossa et al., 2010, Bertossa et al., 2013, Bertossa et al., 2014) and photoperiodic (Saunders 1965) responses, a fully sequenced genome (Werren et al., 2010), and a systemic RNAi response (Lynch and Desplan 2006, Werren et al., 2009). Also of note is Nasonia’s position in the diverse insect order Hymenoptera, an order which evolves more slowly than the order to which *Drosophila* belongs, Diptera (Wyder et al., 2007).

Since the original publication of the *Nasonia* genome(Werren et al., 2010) the assembly has been improved and detailed annotation projects are ongoing (Supplementary Table S1). The level of annotation between assemblies and annotations varies, for example the EvidentialGene dataset (Munoz-Torres et al., 2011) (mapped to the first genome build) contains UTR annotation for 97% of gene models and has a significant amount of GO (gene ontology) annotation, whereas OGS v1.2 (mapped to the latest genome build, adopted by Ensembl) only has UTR annotation for 37% of gene models and has relatively little gene ontology (GO) (Ashburner et al., 2000) annotation. Different gene annotation projects use different sets of gene identifiers, and so comparing studies which use different identifiers to refer to the same genes is not always straightforward. Furthermore, there is currently no existing method for converting identifiers in batch from release to release.

*To whom correspondence should be addressed.

We here present a database combining data from all *Nasonia vitripennis* annotation projects, complete with, for each gene, GO annotations, PFAM domain predictions (Finn et al., 2014), and orthologs in other species. In addition to providing an easy to navigate web interface, WaspAtlas also provides several tools for working with the data including RNAi off-target prediction, GO/PFAM hypergeometric overrepresentation tests, and batch data retrieval for clusters of genes. WaspAtlas is also designed to store and display various types of empirical data, allowing for quick and easy access to published data about each gene regardless of which gene annotation was used in the original study.

## 2 THE DATA

Gene models from four different annotation projects were integrated into WaspAtlas and extant mappings were used to collapse sets of gene identifiers into a single locus where possible (Supplementary Note S1). This process of collapsing gene models into more comprehensive loci also included the merging of associated annotation data (expanded GO annotations, orthologs, empirical data). PFAM protein domain predictions were performed for protein coding genes using HMMer (Finn et al., 2011) (Supplementary Note S2).

For the initial WaspAtlas release, orthologs from 11 different species were calculated using a reciprocal best blast hit (RBH) (Tatusov et al., 1997) approach (Supplementary Note S3), and combined with orthologs calculated by Ensembl (Flicek et al., 2014) in order to facilitate easy comparisons between *N. vitripennis* genes and those of other more well established model organisms. As an example to other authors who wish to include their data in WaspAtlas, two pre-publication datasets have been integrated into the website; a reduced representation bisulfite sequencing (RRBS) dataset describing the differences in gene methylation between wasps exposed to long and short photoperiods (Pegoraro et al, unpublished data), and a conserved non-coding element (CNE) dataset describing the results of a search for highly conserved upstream regulatory elements (Davies et al, unpublished data).

## 3 FEATURES AND USAGE

Access to the WaspAtlas database is provided through a web-based interface. The interface can be conceptually divided into three main components: gene summaries, custom searches, and tools.

The gene summary page for each gene is divided into four sections: i) a brief summary (Fig. 1A), describing the gene identifiers associated with this gene in different annotation releases and their locations on the various genome builds. Also detailed in this section are the annotated GO terms and orthology data, ii) a transcripts section, containing detailed information, illustrations (Fig. 1B), and downloadable sequences for all annotated transcripts, selectable by annotation release, iii) a protein annotation section, containing schematic diagrams of predicted PFAM protein domains and their location within the protein products of each gene (Fig. 1C), again selectable by annotation release, iv) a studies section, containing data from published studies relating to the gene being browsed.

**Fig. 1.**
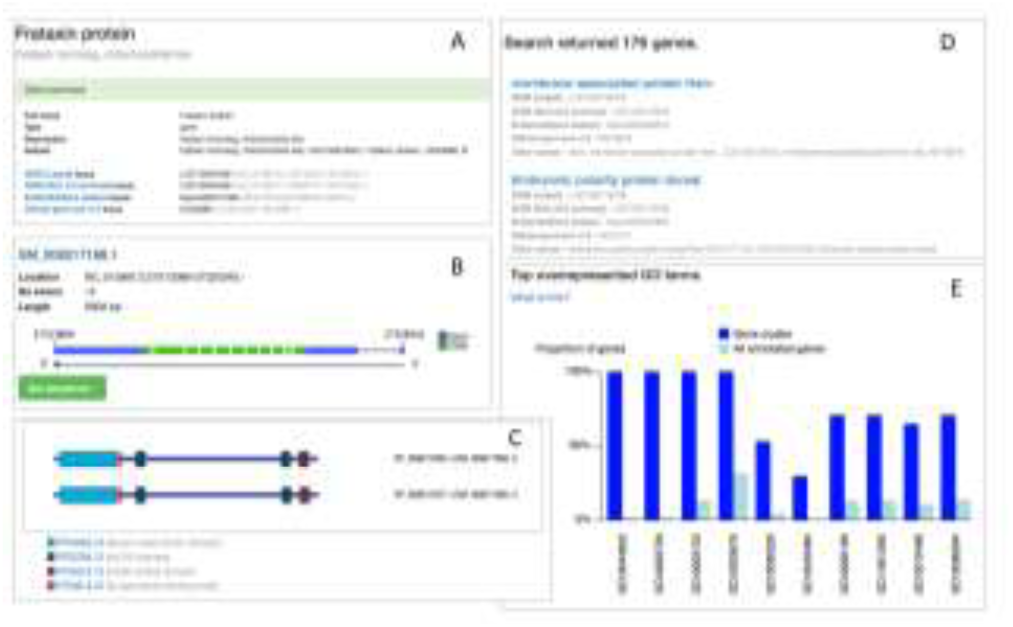
The WaspAtlas database interface. (**A**) Screenshot of part of a brief gene summary. (**B**) Screenshot of a transcript diagram. (**C**) Screenshot of a PFAM protein domain schematic. (**D**) An example of search results for immune response related genes. (**E**) Results for a GO term overrepresentation test on a gene cluster highly enriched for poly-A binding proteins.

Searches on WaspAtlas (Fig. 1D) are accomplished through one of two methods; the quick search or the advanced search. The quick search consists of a single search box located on the right hand side of the navigation bar on every page, and allows a free-text search of gene identifiers, names, descriptions, associated GO terms, and predicted PFAM domains. The advanced search gives the user more control over a search for genes, allowing the user to search using combinations of various gene identifiers/attributes, lists of GO terms and/or PFAM domains, and orthologs in other species.

WaspAtlas tools provide a quick and easy way of performing common analyses with *Nasonia* genes with the latest and most complete functional annotation available. To perform an analysis, the user selects the tool appropriate for their analysis, and fills in the necessary parameters. The user is then given a job identifier which can be used to track the progress of the job. When the job is complete, the user will be given the option to download the raw results as a text file, and depending on the job, given a graphical (Fig. 1E) and/or textual summary of the results. Results of potential interest may also be highlighted; for example the overrepresentation test will highlight the inclusion of large proportions of functional gene families in a given gene cluster. All tools come with detailed explanations of the methods used in the analysis.

## 4 FUTURE DIRECTIONS

A key feature of WaspAtlas that we are keen to expand is its capacity as a central location for published data relating to *Nasonia vitripennis* genes to be viewed. As studies vary widely in their outputs, we are looking to liase with the research community on the best way to present their work within WaspAtlas. In pursuit of this aim, we plan to include more data generated by the community, including RNA-seq expression data and in-situ hybridisation data. The data will be available in tissue specific manner, following the example set by the *Drosophila* database FlyAtlas (Robinson et al., 2013). In addition to the current repertoire of protein-coding genes, we also intend to expand WaspAtlas to cover non-coding RNAs, and pseudogenes, as well as DNA methylation data. In order to further facilitate gene-based analyses, we will increase the selection of available tools to include more commonly performed analyses, and intend to expand the annotation of orthologs in other species in the near future.

